# Signatures of genetic variation in human microRNAs point to processes of positive selection related to population-specific disease risks

**DOI:** 10.1101/2021.05.24.445417

**Authors:** Pablo Villegas-Mirón, Alicia Gallego, Jaume Bertranpetit, Hafid Laayouni, Yolanda Espinosa-Parrilla

## Abstract

The occurrence of natural variation in human microRNAs has been the focus of numerous studies during the last twenty years. Most of them have been dedicated to study the role of specific mutations in diseases, like cancer, while a minor fraction seek to analyse the diversity profiles of microRNAs in the genomes of human populations. In the present study we analyse the latest human microRNA annotations in the light of the most updated catalog of genetic variation provided by the 1000 Genomes Project. We show by means of the *in silico* analysis of noncoding variation of microRNAs that the level of evolutionary constraint of these sequences is governed by the interplay of different factors, like their evolutionary age or the genomic location where they emerged. The role of mutations in the shaping of microRNA-driven regulatory interactions is emphasized with the acknowledgement that, while the whole microRNA sequence is highly conserved, the seed region shows a pattern of higher genetic diversity that appears to be caused by the dramatic frequency shifts of a fraction of human microRNAs. We highlight the participation of these microRNAs in population-specific processes by identifying that not only the seed, but also the loop, are particularly differentiated regions among human populations. The quantitative computational comparison of signatures of population differentiation showed that candidate microRNAs with the largest differences are enriched in variants implicated in gene expression levels (eQTLs), selective sweeps and pathological processes. We explore the implication of these evolutionary-driven microRNAs and their SNPs in human diseases, such as different types of cancer, and discuss their role in population-specific disease risk.

## Introduction

MicroRNAs (miRNAs) are short (~22 nucleotides) single-stranded regulatory non-protein-coding RNAs that perform a post-transcriptional negative control of the expression of more than 60% of the whole human genome (Friedman et al. 2009). They are involved in the control of almost every cellular process, including development, differentiation, proliferation and apoptosis, and present important roles in diseases. They are transcribed by RNA polymerase II as primary sequences, which are later processed by the proteins Drosha and Dicer into a miRNA duplex formed by two mature miRNA strands, 5p and 3p (Ha et al. 2014). This mature molecule is then loaded onto an AGO protein forming the RNA-induced silencing complex (RISC), promoting the RNA silencing by translation repression or mRNA degradation. Target gene repression is accomplished by the partial sequence complementarity between the target mRNA and the miRNA. In this interaction, a perfect match between the miRNA seed region, expanded across nucleotides 2-8 of the 5’ extreme, and the target site, usually located within the mRNA 3’ untranslated region, is needed (Lewis et al. 2005; Grimson et al. 2007; Bartel et al. 2009; Berezikov 2011). Other positions of the mature sequence may interfere in the mRNA binding, like the 3’ supplementary and compensatory sites, that enhance the seed-matched binding efficiency (Grimson et al. 2007; Friedman et al. 2009; Bartel 2018).

miRNAs have experienced multiple periods of fast turn over and lineage-specific expansions through their evolutionary trajectory (Lu et al. 2008; Iwama et al. 2012). Most of the current human miRNAs originated in two accelerated peaks of miRNA expansion that are reported during mammalian evolution: the first peak of new miRNAs was located at the initial phase of the placental radiation, while the second and highest peak was observed at the beginning of the simian lineage, that originated more than a half of the current repertoire (Iwama et al. 2012; Santpere et al. 2016). These miRNA expansions were implicated in the acquisition of new regulatory tools that have been directly linked with animal complexity and evolutionary innovations across all lineages (Hertel et al. 2006; Heimberg et al. 2008; Wheeler et al. 2009).

miRNAs can be found either in intergenic regions or being hosted by other elements, like protein-coding and non-coding genes or repetitive elements like transposons. These are the genomic contexts where hairpin-like transcripts initially emerge and are gradually shaped by evolution until they become functional miRNAs (Berezikov 2011). Differences in the genomic environment and location of miRNAs are associated with different evolutionary properties. For example, in França et al. 2016 the authors show the association of the age of the host gene with the breadth expression and evolutionary trajectory of recently emerged hosted miRNAs. Duplication events are one of the main sources of new miRNAs. These can be found close to each other when the duplication is local, forming clusters that are found to be evolutionary related and functionally implicated in similar regulatory pathways (Wang et al. 2016). The origin of miRNAs and their target sites are tightly related to the dynamics of transposable elements (TE). These are sequences that jump, replicate and insert in other parts of the genome, generating mutations. However, apart from the damaging consequences of these changes, they can also incorporate new functional regions in other genomic environments (like miRNA target sites) and modify regulatory networks (Feschotte 2008; Chuong et al. 2017). According to some authors (Piriyapongsa et al. 2007; Qin et al. 2015; Petri et al. 2019), the expansion of new miRNAs in the primate lineage gave birth to a great number of TE-derived miRNAs, highlighting the importance of transposons as a source of genomic innovation.

The computational analysis of human genetic variation has traditionally been focused on protein-coding genes, being non-protein coding sequences neglected from this kind of studies. However, in recent years, several reports have paid more attention to the consequences of naturally occurring variation in miRNAs (Cammaerts et al. 2015). A signature of purifying selection shapes the miRNA diversity worldwide, revealing that human miRNAs are highly conserved sequences that rarely accept changes in their sequences (Quach et al. 2009), indeed miRNA expression and functionality are usually tightly subjected to the presence of variants within (Quach et al. 2009) and outside (Borel et al. 2011) their hairpin. Sequence changes in the premature and loop regions might generate distorsions in their folding and affect the expression and maturation of the primary sequences (Fernandez et al. 2017). Moreover, the occurrence of changes in the mature region and the seed (Gong et al. 2012; Hill et al. 2014; Gallego et al. 2016; He et al. 2018), which outstands as the most conserved region of the hairpin, can dramatically affect the recognition of their target genes, which is also affected by the presence of variants in their target sites (Li et al. 2012). All these changes might induce massive rewirings of the miRNA regulatory networks and alter the downstream processes, inducing gene expression changes and phenotypic variation that might degenerate in pathogenic processes (Sethupathy and Collins 2008; Rawlings-Goss et al. 2014; Ghanbari et al. 2017; Grigelioniene et al. 2019), but also be the origin of genetic innovations responsible for phenotypic adaptations (Lu et al. 2012). Several authors have reported population-specific variants that affect different dimensions of the miRNA functionality (Saunders et al. 2007; Torruella-Loran et al. 2016) and their target sites (Li et al. 2012) and might be involved in adaptation processes. More recently, it has been reported a clear signal of adaptive evolution in a metabolic-related miRNA responsible for adaptations to past famine periods (Wang et al. 2020).

In this study we revisited the hosting and conservation patterns of the most complete human miRNA catalog to date. We also performed a comprehensive computational analysis of their diversity patterns worldwide, considering the factors that might contribute the most to the configuration of this variation. We finally studied the population differences and putative positive selection signals of the variable miRNAs, and looked at the potential consequences of this variation in terms of human diseases and recent adaptation.

## Results

### The genomic context of miRNAs is associated with their evolutionary age

To study the recent evolutionary history of human miRNA genes a total of 1918 precursor miRNAs (miRBase v.22, March 2018) were considered, from which 1904 remained after *liftOver* conversion to the hg19 assembly (Supplementary Table S1). From these miRNA precursors, 50.3% presented a complete annotation of their mature sequences (5p and 3p), while the other half presented only a single mature sequence identified in one of their arms (Fig. 1a and Supplementary Fig. S1). First, we classified these 1904 miRNAs in groups of conservation, according to their evolutionary age, by adapting the categories from Iwama et al. (2013) and Santpere et al. (2016) (see Methods). In total, 1623 (85.2%) miRNAs were classified in four different conservation categories: Primates (985, 51.7%), Eutherians (421, 22.1%), Metatheria-Prototheria (63, 3.3%) and conserved beyond mammals (154, 8%). The remaining miRNAs (281, 14.8%) could not be classified due to the absence of data or discrepancies between studies and were excluded from the subsequent analyses (Supplementary Table S1).

**Fig. 1.**
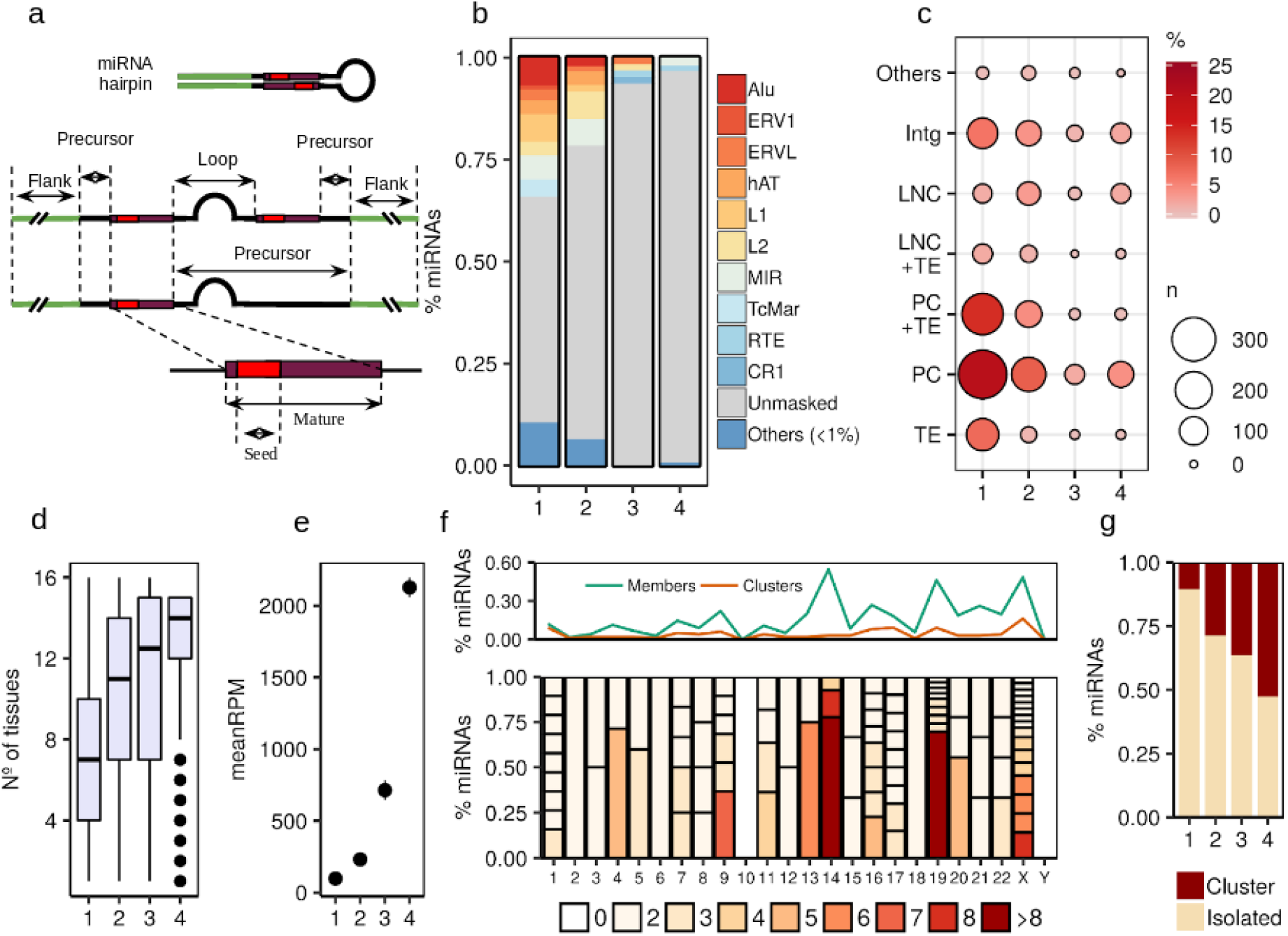
Description of human miRNAs in terms of genomic context, evolutionary age groups (see Methods), expression levels and clustering. **(a)** Description of the miRNA hairpin regions identified and analysed in the study. Not all the primary sequences present two mature sequences annotated by miRbase. When the two mature sequences are not given (incomplete annotation), the precursor region is extended from the first mature to the other flanking region. **(b)** TE-derived miRNA frequencies across conservation groups (Primates, 1; Eutherians, 2; Metatheria and Prototheria, 3; Conserved beyond mammals, 4). **(c)** Integrated hosting of miRNAs that shows the combination of the different hosting elements that overlap with miRNA sequences. The “Other” group is made with the minor categories (PC+LNC and PC+LNC+TE) that represent less than 1% of the total dataset (Supplementary Table S2). **(d)** Number of tissues where the miRNA is expressed across evolutionary ages **(e)** Mean expression level (Reads per million mapped reads; RPM) of miRNAs across evolutionary ages **(f)** Whole genome clustering patterns of miRNAs. The upper plot represents the frequency of miRNAs that belong to a certain cluster in each chromosome (Members) and the frequency of clusters in the whole genome (Clusters). The lower plot represents the miRNA clusters per chromosome, according to the number of members and their frequency among the clustered miRNAs. **(g)** Fraction of clustered and isolated miRNAs across evolutionary ages

Next, we classified miRNAs in different genomic contexts by identifying the different elements that overlap their precursor sequences. According to GENCODE 19 (v.29) we found that 483 (25%) miRNAs fell in intergenic regions (Intg), while 1421 were located within protein coding genes (PC) (1217, 63.9%) and long non-coding RNAs (LNC) (204, 10.7%), either presenting a single or multiple overlapping host genes. In our dataset we found that 856 (60%) intragenic miRNAs (protein coding and lncRNA) overlapped introns of the host sequence, while 545 (38%) were located within exonic regions. The remaining 20 (~1%) showed a mixture of intronic/exonic locations (Supplementary Table S1). Further, we used the last release of the RepeatMasker database (Smit et al. 2013-2015) to identify the different forms of transposable elements (TEs) and repetitive sequences that host miRNAs. We found 660 (35%) miRNAs overlapping TEs alone or in combination with other genes, while the remaining 1244 (65%) were either unmasked or overlapping other forms of repetitive sequences and genes. Interestingly, we found a strong correlation between the frequencies of the TE-hosting miRNAs and their evolutionary age, being the primate-specific group the one with the highest presence of miRNAs in this context (440, 23.1%; Fig. 1b). Alu (67, 6.8%), L1 (54, 5.4%), TcMar (42, 4.2%) and the LTR elements ERV1 and ERVL (36, 3.6%) were found mainly among the primate-specific miRNAs, while hAT (3.3%) and L2 (28, 6.6%) elements were also present in the eutherian group (Supplementary Table S2). It is of interest to note that the contribution of MIR (15.3%) and DNA elements like TcMar (14.8%) and hAT (12.8%) families to the miRNA context is higher than to the whole genome (Supplementary Fig. S2a).

We found that the genomic context increased in complexity when different elements appeared hosting the same miRNA simultaneously. We studied the integrated hosting of miRNAs across the conservation groups considering the different combinations of elements (Fig. 1c, Supplementary Table S3). This shared hosting evidences the two main sources of miRNAs: protein coding genes (796; 41.8%) and TEs (196; 10.2%), with 401 miRNAs presenting a combination of both (21%). As expected, the genomic context is associated with the age of miRNAs (Chi square test = 238.25, p = 2.2e-16). This association shows that primate-specific miRNAs present a dominance of overlapping TEs in comparison with non-primate miRNAs, with the TE and TE + PC hosting categories being the major contributors across environments. On the other hand, lncRNAs are highly associated with the miRNA context among the non-primate groups, mainly in the group of miRNAs conserved beyond mammals (Supplementary Fig. S2b).

We made use of the miRNA expression levels in 16 different human tissues extracted from Panwar et al. (2017) (see Methods) to study their correlation across groups of conservation. As seen in Fig. 1d, the tissue specificity is higher at lower evolutionary ages, which indicates the limited expression breadth of young miRNAs. Also, the expression levels were correlated with age, having the more conserved miRNAs an overall higher expression due to their consolidated role in regulatory networks (Fig. 1e).

Due to the evolutionary relevance of the miRNA organization in the genome, we revisited the clustering patterns of the miRBase annotations. When studying the closeness between miRNAs, an increment of distances ranging 1-10kb was found (Supplementary Fig. S2c), which indicates a high accumulation of close miRNAs in certain regions. According to this, we defined that two miRNAs belong to the same cluster when they are located 10kb or closer from each other. A total of 100 clusters were identified in the whole genome (Fig. 1f and Supplementary Fig. S3), represented by 352 miRNA members. Two thirds of these clusters (64) were constituted only by two genes, while 36 clusters presented more than two. Two main clustering hotspots were observed in the chromosomes 14 (42) and 19 (46), as previously reported by Guo et al. (2014), while the X chromosome presented a similar amount of clustered miRNAs (57) but more widespread in different smaller groups (Fig. 1f). A total of 1552 miRNAs were located in isolated regions. We also found a strong correlation between the clustering patterns of miRNAs and groups of conservation (Fig. 1g). The more conserved miRNAs tend to be found in clusters rather than in isolated regions, something likely related to the conserved role of clustered miRNAs in similar biological processes (Berezikov 2011; Wang et al. 2016).

### Nucleotide diversity of miRNAs is strongly shaped by their age, genomic context and localization

The genetic variation of the miRNA dataset was analysed in the different miRNA functional regions using human genetic variation from the 1000 Genomes project (Fig. 1a; Auton A et al. 2015). A total of 569 single nucleotide polymorphisms (SNPs) were located in 466 miRNA precursors (26.1%), but when considering a region of the same size at both sides of the precursor sequence (5’ and 3’ flanking regions) the number of SNPs increased to 1994 in 1026 miRNAs (55.9%). Therefore, more than half of the variability found in our miRNAs comes from the neutral-like flanking regions. The mature sequence is considered the most conserved and important functional region of the miRNA, since it regulates the target gene by binding to the 3’UTR mainly through the seed region. In our dataset, 212 SNPs were present in 194 mature sequences (7.5%), while 79 SNPs were present only in the seed region of 75 miRNAs (2.9%).

To study the sequence variation of human miRNAs we analysed the nucleotide diversity of 1904 miRNA precursor sequences described in miRBase in the pooled population sample from the 1000 Genomes project. The genomic context refers to the environment where miRNAs originally emerged, which might be determinant to their level of variation. We calculated the global nucleotide diversity (Pi) in the whole precursor sequence by considering the age, location and clustering of the miRNAs (Fig. 2). We found significant differences when comparing the Pi of miRNAs in the different contexts (Kruskal-Wallis p = 0.013). Fig. 2a shows that miRNAs harboured by TEs exhibit a significantly higher Pi than in other genomic contexts. Next, we examined the TE-family specific diversity of the hosted miRNAs and wondered which TE families contribute more to this high diversity (Supplementary Fig. S3a). We performed a multiple linear regression analysis with the different families as predictors and found that Alu and ERVL are significantly associated with the increase of nucleotide diversity (Alu, p = 0.013; ERVL, p = 5.11e-04).

**Fig. 2.**
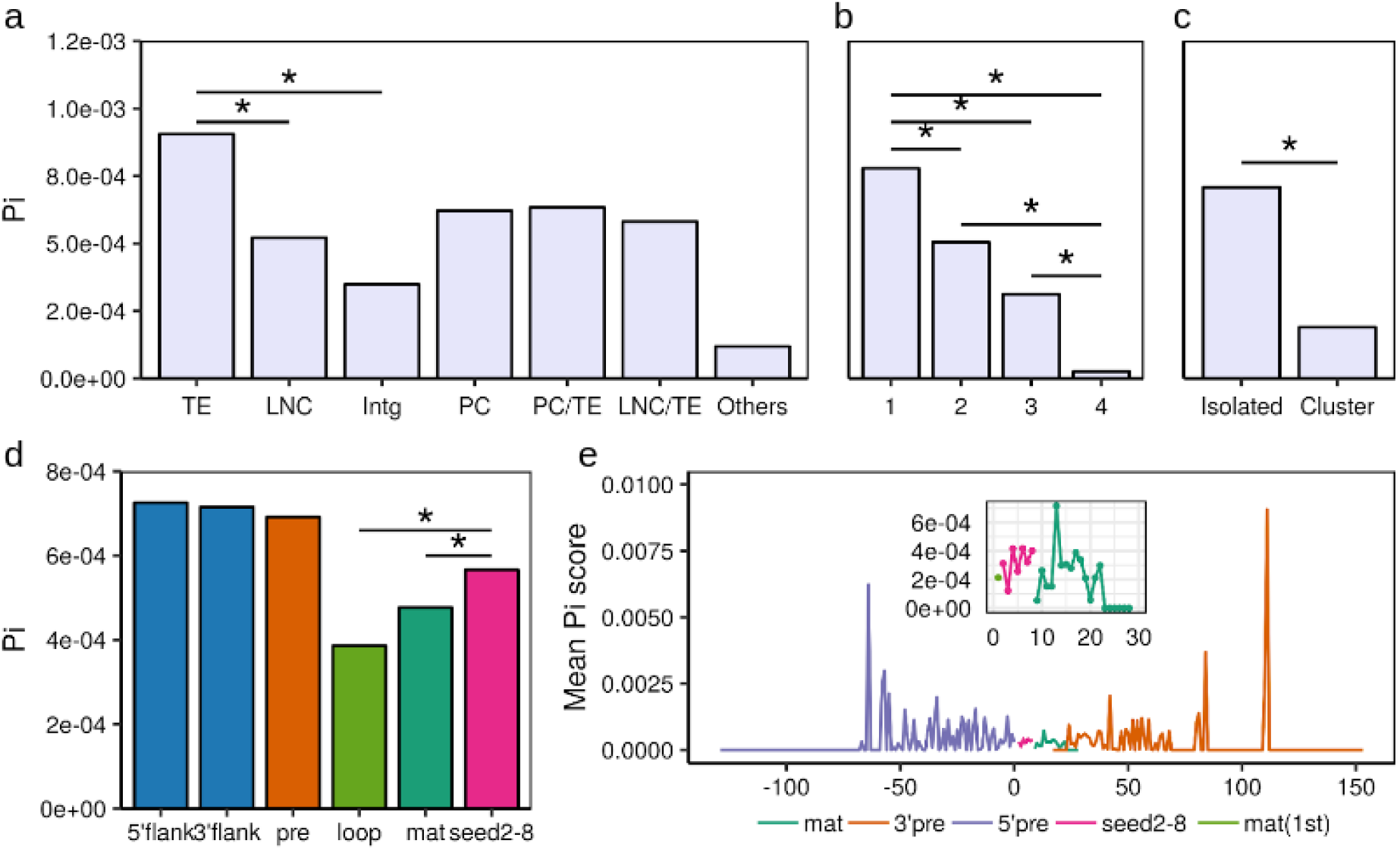
Mean nucleotide diversity differences between miRNAs in different annotation categories and functional regions. **(a)** Differences between the genomic contexts where the human miRNAs are found. Wilcoxon pairwise comparisons (Bonferroni corrected) show that transposable elements (TE) present a significantly higher diversity than other environments (TE vs LNC, p = 0.022; TE vs Intg, p = 0.022. **(b)** Differences across miRNA conservation groups (see Methods). Primate-specific miRNAs (group 1) show a significantly higher diversity in comparison with the others (1 vs 2, p = 0.00057; 1 vs 3, p = 0.0178; 1 vs 4, p = 3.93e.10; Wilcoxon pairwise comparisons, Bonferroni corrected). Significant differences are also seen for the miRNAs conserved beyond mammals (group 4) (4 vs 3, p = 0.0178; 4 vs 2, p = 2.6e-05; Wilcoxon pairwise comparisons, Bonferroni corrected). **(c)** Differences between miRNAs found isolated and organised in clusters. Isolated miRNAs are associated with a significantly higher diversity than the members of clusters (Wilcoxon test, p = 3.663e-10). **(d)** Diversity comparison between the different functional regions identified in the miRNA hairpins. The seed region (2-8 nucleotides) presents a significantly higher diversity than other conserved regions (seed vs loop, p = 0.0011 and seed vs mat, p = 0.0056; Wilcoxon pairwise comparisons, Bonferroni corrected). **(e)** Mean nucleotide diversity calculated in each relative position of the precursor miRNA. The zoomed region correspond to the diversity per position found in the mature sequence

As expected, the evolutionary age is another determinant factor in the miRNA sequence diversity. We found that Pi presents a clear correlation with the miRNA conservation (Fig. 2b; see Methods), with significant differences among the different groups (Kruskal-Wallis p = 2.373e-11). The highest diversity was seen in the miRNAs classified as primate specific (group 1) and the lowest in those conserved beyond mammals (group 4).

Regarding the clustering patterns of miRNAs, we found that diversity differences between clustered and isolated miRNAs reached significant levels (Wilcoxon p = 3.663e-10) (Fig. 2c) which, as seen before, it might be a reflection of the higher conservation of clusters due to their functionality in cooperative processes (Wang et al. 2016; Kabekkodu et al. 2018) and also the fact that most of the clustered miRNAs have originated after common duplication events (Hertel et al. 2006).

Considering the above, sequence diversity levels of human miRNAs seem to be driven by their location, age and genomic context. These factors might also determine the presence of mutations in miRNA sequences that could affect their expression, hairpin folding and even their ability to bind their target genes and, therefore, be determinant for their evolutionary trajectory. Because of that, we wanted to study the integrated contribution of these factors to the observed diversity differences. We applied a multiple linear regression model to the diversity data and the different miRNA categories (genomic context, evolutionary age and clustering). The regression model showed that age (being primate specific, p = 3.3e-03), clustering (being isolated, p = 3.6e-04) and genomic context (not being intergenic, p = 0.015) are predictors significantly associated with the increase of Pi in human miRNAs.

### An excess of diversity in the seed region is driven by a reduced number of miRNAs

The analysis of the nucleotide diversity (Pi) across different miRNA regions indicated an overall higher diversity in the precursor and flanking regions compared to the rest of regions (Wilcoxon test p < 0.05). Surprisingly the loop region presented the lowest diversity of the whole miRNA hairpin (Fig. 2d). This might reflect the importance of this region in the hairpin folding, which is determinant for the processing of the primary sequence. Previous studies (Torruella-Loran et al. 2016) showed that the seed is the most conserved region of the miRNA, which has been associated with its functional relevance due its central role in target binding. However, our results showed a higher Pi in the seed than in other conserved regions, like the mature (outside seed) and the loop (Wilcoxon pairwise comparisons p = 0.0011 and p = 0.0056, respectively). It is worth noting that this level of diversity in the seed comes from the variation of a small set of miRNAs (75, 2.9%), showing that, indeed, most of the human miRNAs are conserved in their seed. On the other hand, the seed region presented values of SNP density similar to those in the mature outside the seed (Supplementary Fig. S4b), which suggests that, considering the values of nucleotide diversity, the seed region is more populated by high frequency variants than the mature region. The region-specific levels of diversity were studied in the whole range of minor allele frequency (MAF), where the seed region was consistently found with diversity levels below the mature region until a frequency ~ 50% (Supplementary Fig. S4c). This shows that no bias in the variant content is confounding these results. Overall, these data suggest that the high diversity observed in this set of miRNAs might be a consequence of the specific targeting of positive selection processes, as discussed below.

Previous reports on miRNA targeting (Grimson et al. 2007; Wheeler et al. 2009) show that not only the seed region but also certain positions in the mature sequence are involved in target binding. To further analyse the variation in the miRNAs, nucleotide diversity was studied at position basis in the whole precursor sequence (Fig. 2e). As expected, the general pattern shows that the mature sequences are located in a valley of diversity, which confirms their overall conservation. Different levels of diversity are seen in the mature sequence. More specifically a decrease in diversity is seen at the 3’ end, corresponding to the region known as participating in the complementary binding of mRNAs.

### Highly differentiated miRNA SNPs are enriched in signals of positive selection and expression variation

The excess of diversity found in the seed region may respond to particular processes of positive selection that generate frequency shifts at population level. These population-specific changes could affect the miRNA binding to the target gene and change the targeting profiles.

In this line, we wanted to study the population-specific patterns of diversity found within the miRNA seed regions. In Supplementary Fig. S5 we show the Pi values of the seed regions from a total of 60 miRNAs presenting genetic variants (DAF >= 5%) calculated in each of the 26 populations of our study. The clustering pattern of diversity sharing among populations reflects the similarities of demographic and potential evolutionary histories in the same continental group. As expected, African populations (AFR) are clustered separately from the other populations, showing the highest differentiation probably due to the Out-of-Africa event. A higher diversity sharing is seen among the non-African populations. There are some clear continental-specific groups of miRNAs that might be the result of demographic dynamics and/or genetic drift, but also of local processes of positive selection on certain alleles. Considering the group-specific membership of miRNA alleles we found that 37% (22) are exclusively present in AFR, while 13% (8) are found in non-Africans, private or shared among other groups (European (EUR), American (AMR), EastAsian (EAS) and South Asian (SAS)). The other alleles are shared between African and non-African populations (50%, 30), being 21 (35%) present in all continents.

Next, mean population differentiation (F_st_) values across all possible population comparisons were calculated for the different miRNA regions (Fig. 3a). As shown, the seed presents an overall F_st_ score higher than the rest of the mature sequence in almost all the compared groups. This tendency is stronger in comparisons including AFR populations than non-African ones. Although demographic dynamics are generally the main cause in the existing differentiation between populations, the high F_st_ values in the seed, compared to other conserved regions like the mature (outside seed) and the loop, suggest that this region could have been particularly targeted by processes of positive selection. Surprisingly, in contrast with the overall low diversity values seen before, the loop region also exhibits particularly high F_st_ scores in some comparisons, especially in the AFR vs SAS populations.

**Fig. 3.**
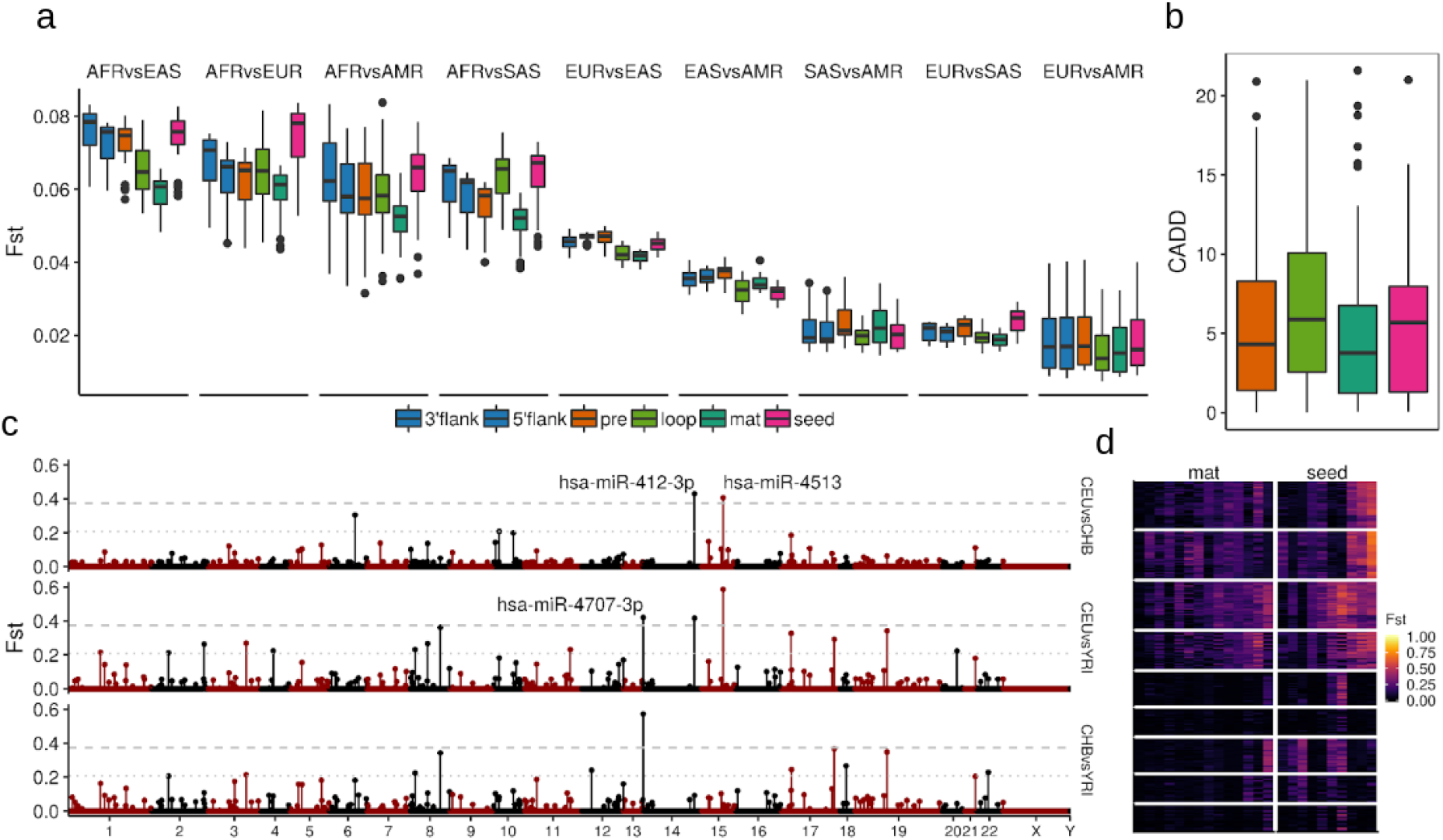
Analysis of F_st_ values across miRNA regions and candidates. **(a)** Mean F_st_ values per miRNA region across all population comparison groups. The F_st_ values were calculated in all the variant regions. **(b)** Combined Annotation Dependent Depletion (CADD) scores distributions, as a measure of the predicted level of deleteriousness of the variants, across miRNA regions **(c)** Manhattan plot showing the mean F_st_ values per miRNA mature sequence in the three comparisons of reference. Two F_st_ thresholds were used to extract the potential miRNA candidates under positive selection (1% and 5%). **(d)** Heatmap showing the per-SNP F_st_ values of the variants found in the mature outside seed (14) and seed (10) region of the top 5% miRNA candidates, where the columns correspond to SNPs and rows to all possible population comparisons (243)

Further, we evaluated the potential functionality of the precursor region-specific SNPs by contrasting their overall Combined Annotation Dependent Depletion (CADD) score distributions, a statistic designed to measure the deleteriousness of human variants (Rentzsch et al. 2019). As shown in Fig. 3b, the CADD scores associated with the loop and seed regions are slightly higher than the rest of the precursor sequence, although non-significant. This evidence reinforces the idea that these regions are specifically implicated in processes potentially involved in adaptive selection.

We wanted to examine the extent to which the top F_st_ scoring SNPs participate in putative signatures of recent positive selection. We focused on signals characterized by the presence of long haplotypes at high (ongoing hard sweeps) and moderate frequencies (soft sweeps) in individual populations, detected by the statistics integrated haplotype score (iHS) (Voight et al. 2006) and the number of segregating sites by length (nSL) (Ferrer-Admetlla et al. 2014) (see Methods). We pooled the SNP set (100, 16%) that showed extreme F_st_ values (>99%) in the whole miRNA precursor sequence in all population comparisons, and explored their involvement in selective sweeps. Among these top SNPs we found that 23% and 18% present extreme iHS and nSL scores (≥ 2), respectively, in at least one population, while the proportion of highly scoring SNPs in the whole dataset is only 13.8% (iHS) and 11.5% (nSL). This result suggests that highly differentiated SNPs in the precursor miRNA sequence are more likely to be found in genomic regions that hold signatures consistent with recent positive selection signatures (iHS Chi square test = 11.29, p = 7.77e-04; nSL Chi square test = 6.74, p = 9.38e-03).

Nucleotide changes in regions involved in miRNA sequence processing (pre, loop) and target binding (mature, seed) might affect the regulation of their target genes and, therefore, generate expression variation that could lead to genetic disorders, but also to phenotypic adaptations. We used the Genotype-Tissue Expression (GTEx) Project catalog (v7) of associated eQTL-eGene pairs to study the potential impact of our miRNA-harbouring top SNPs in gene expression variation (Aguet F et al. 2017). Among the top 100 SNPs in the precursor sequences, 54% (54) are reported as significant expression Quantitative Trait Loci (eQTLs) by GTEx, while the 24.7% (154) are found in the whole SNP dataset. Also, we used the most recent release of the genome-wide association studies (GWAS) catalog (v1.0) (Buniello et al. 2019) to evaluate the extent to which these highly differentiated SNPs are associated with genetic diseases and traits. In this case, 5% (5) of the top SNPs present significant associations in GWAS studies, while only 1.7% (11) are found in the whole SNP dataset. These results indicate that highly differentiated miRNA-harbouring SNPs are more likely to be reported as significant eQTLs (Chi-square test = 33.994, p = 5.528e-09) and GWAS associated SNPs (Chi square test = 6.7841, p = 9.19e-03), which suggests their implication in expression variation and human diseases.

### miRNA recent evolution might be driven by targeted processes in their seed related to positive selection and disease

In order to identify potential miRNA candidates under the selection pressures of local adaptations, we calculated mean F_st_ values in the whole mature sequence. Fig. 3c shows the genome wide distribution of mature-specific F_st_ values in the three comparisons of reference (Utah Europeans (CEU) vs Han Chinese (CHB), CEU vs Yoruba (YRI) and CHB vs YRI), where three miRNAs are found in the top 1% (hsa-miR-1269b, hsa-miR-412-3p, hsa-miR-4707-3p) and 22 above the 5% (Table 1). Surprisingly the three most divergent miRNAs belong to conservation groups older than primate specific, which suggests that these population-specific changes might respond to potential adaptations that affect well-established regulatory pathways. These candidate miRNAs harbour 10 SNPs within their seed regions (10 miRNAs) and 14 SNPs in other positions of the mature sequence (14 miRNAs). As seen in Fig. 3d, seed-harboring SNPs like rs2273626 (hsa-miR-4707-3p) present the most extreme F_st_ scores in the candidate mature sequences and reach top values (>99.98%) in the whole miRNA distribution. Among these, seven SNPs in both seed (rs6771809, rs77651740, rs28655823, rs2273626, rs2168518, rs7210937, rs3745198) and mature regions (rs56790095, rs73239138, rs404337, rs2155248, rs61992671, rs12451747, rs73410309) were reported by GTEx as significantly associated to gene expression variation.

**Table 1.**
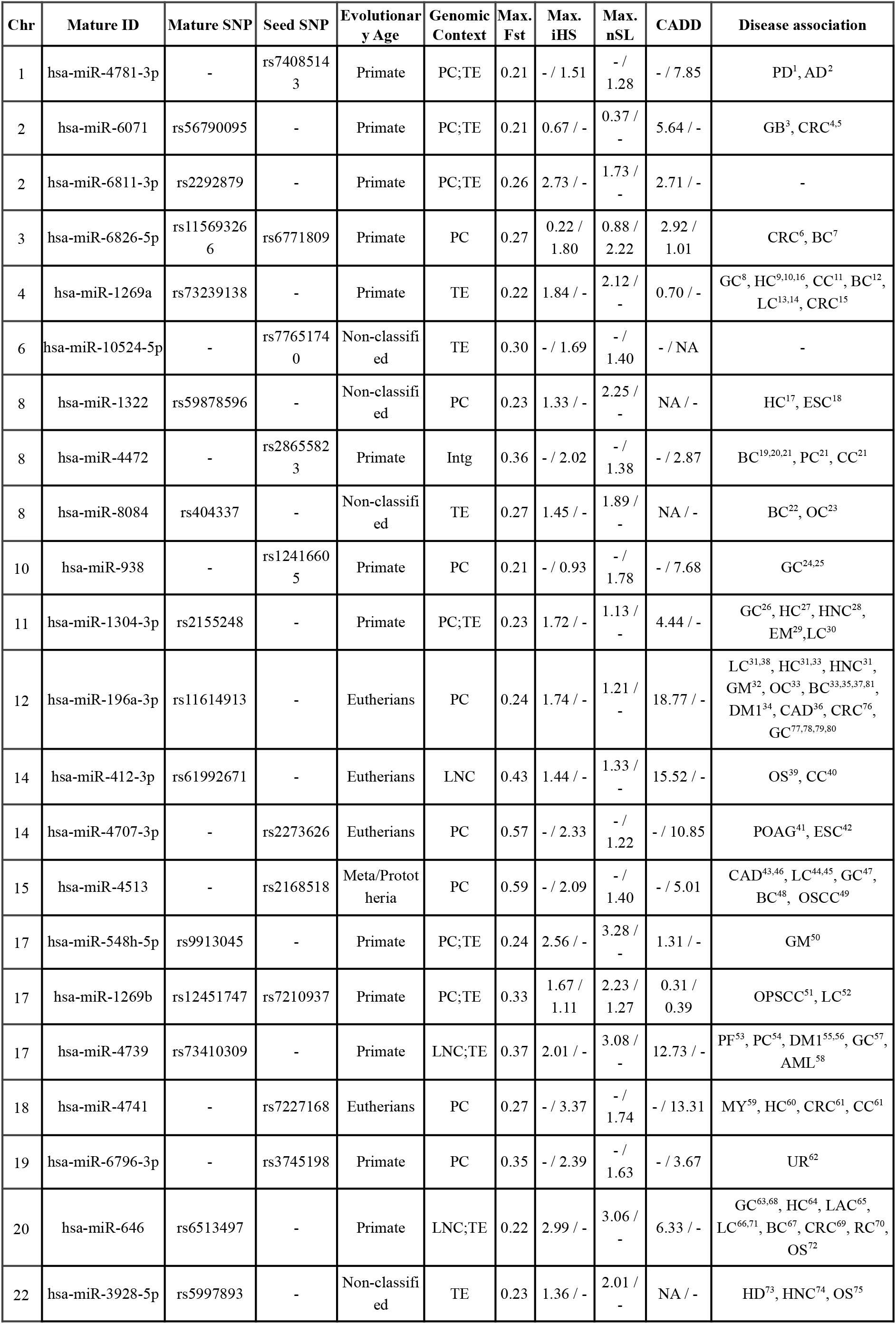
Top 5% miRNA candidates of different ages and genomic contexts under putative positive selection. The Max. Fst value represents the maximum mean Fst of the mature sequence among the three comparisons of reference. The selection test values (iHS and nSL) correspond to the population that exhibit the maximum value of the mature (left) and seed SNP (right). The CADD column provides the predicted deleteriousness scores (see Methods) of the mature (left) and seed SNP (right). Disease association for most of the candidates are indicated in the disease column and some examples are described in the main text: AD (Alzheimer’s disease; (2) Satoh et al. 2015), AML (acute myeloid leukemia; (58) Cattaneo et al. 2015), BC (breast cancer; (7) Danková et al. 2020, (12) Sarabandi et al. 2021, (19) Li et al. 2020, (20) Wang et al. 2018, (21) Kim et al. 2012, (22) Gao et al. 2018, (33) Choupani et al. 2019, (35) Ahmad and Shah 2020, (37) Zhao et al. 2016, (48) Li et al. 2019, (67) Darvishi et al. 2020), CAD (coronary artery disease; (36) Fragoso et al. 2019, (43) Mir et al. 2019, (46) Li et al. 2015), CC (colon cancer; (11) Mao et al. 2017, (21) Kim et al. 2012, (40) Zhu et al. 2020), CRC (colorectal cancer; (4,5) Slattery et al. 2018, (6) Kijima et al. 2017; (15) Bu et al. 2015, (61) Cojocneanu et al. 2020, (69) Dai et al. 2017), DM1 (type 1 diabetes mellitus; (34) Ibrahim et al. 2019, (55) Delić et al. 2016, (56) Li et al. 2018), EM (endometriosis; (29) Xu et al. 2017), ESC (esophageal squamous cell carcinoma; (18) Zhang et al. 2013, (42) Bi et al. 2020), GB (glioblastoma; (3) Zhou et al. 2020), GC (gastric cancer; (8) Li et al. 2017, (24) Torruella‐Loran et al. 2019, (25) Arisawa et al. 2012, (26) Kurata and Lin 2018, (47) Ding et al. 2019, (57) Dong et al. 2015, (63) Cai et al. 2016, (68) Zhang et al. 2017), GM (glioma; (32) Yang et al 2020, (50) Ji et al. 2020), HC (hepatocellular carcinoma; (9) Min et al. 2017, (10) Xiong et al. 2015, (16) Wang et al. 2019, (17) Zhao et al. 2020, (27) Oura et al. 2019, (60) Liu et al. 2019, (64) Wang et al. 2014), HD (Huntington disease; (73) Reed et al. 2018), HNC (head and neck squamous cell carcinoma; (28) Petronacci et al. 2020, (74) Fadhil et al. 2020), LAC (laryngeal carcinoma; (65) Yuan et al. 2020), LC (lung cancer; (13) Jin et al. 2018, (14) Wang et al. 2020; (30) Othman et al. 2013, (31) Liu et al. 2018, (38) Wang et al. 2017, (44) Ghanbari M et al. 2014, (45) Ghanbari M et al. 2017, (52) Yang et al. 2020, (66) Wang et al. 2020, (71) Pan et al. 2016), MY (myeloma; (59) Zhang et al. 2019), OC (ovarian cancer; (23) Chong et al. 2015, (33) Choupani et al. 2019), OPSCC (oral and pharyngeal squamous carcinoma; (51) Chen et al. 2016), OS (osteosarcoma; (39) Martin-Guerrero et al. 2018, (72) Sun et al. 2015, (75) Xu et al. 2014), OSCC (oral squamous cell carcinoma; (49) Xu et al. 2019), PC (prostate cancer; (21) Kim et al. 2012, (54) Wang et al. 2020), PD (Parkinson disease; (1) Beecham et al. 2015), PF (pleural fibrosis; (53) Wang et al. 2019), POAG (open-angle glaucoma; (41) Ghanbari, et al. 2017), RC (renal carcinoma; (70) Li et al. 2014), UR (urolithiasis; (62) Liang et al. 2019).

As seen before, the presence of SNPs in the seed region might lead to variations of the miRNA targeting profiles. In order to evaluate the degree of change that a single SNP might generate, we adapted the *TargetScanHuman* (Agarwal et al. 2015) pipeline to predict the allele-specific targets of the seed-variant candidates. When comparing the sets of target genes due to the ancestral and derived alleles we observed that, among the top ten miRNAs with SNPs in their seed, only two present a cosine similarity (see Methods) above 70% (hsa-miR-10524-5p and hsa-miR-4513), while the other candidates fall below 23%. This indicates the dramatic target shift that a single SNP generates and might be involved in regulatory adaptations (Table 2).

**Table 2.**
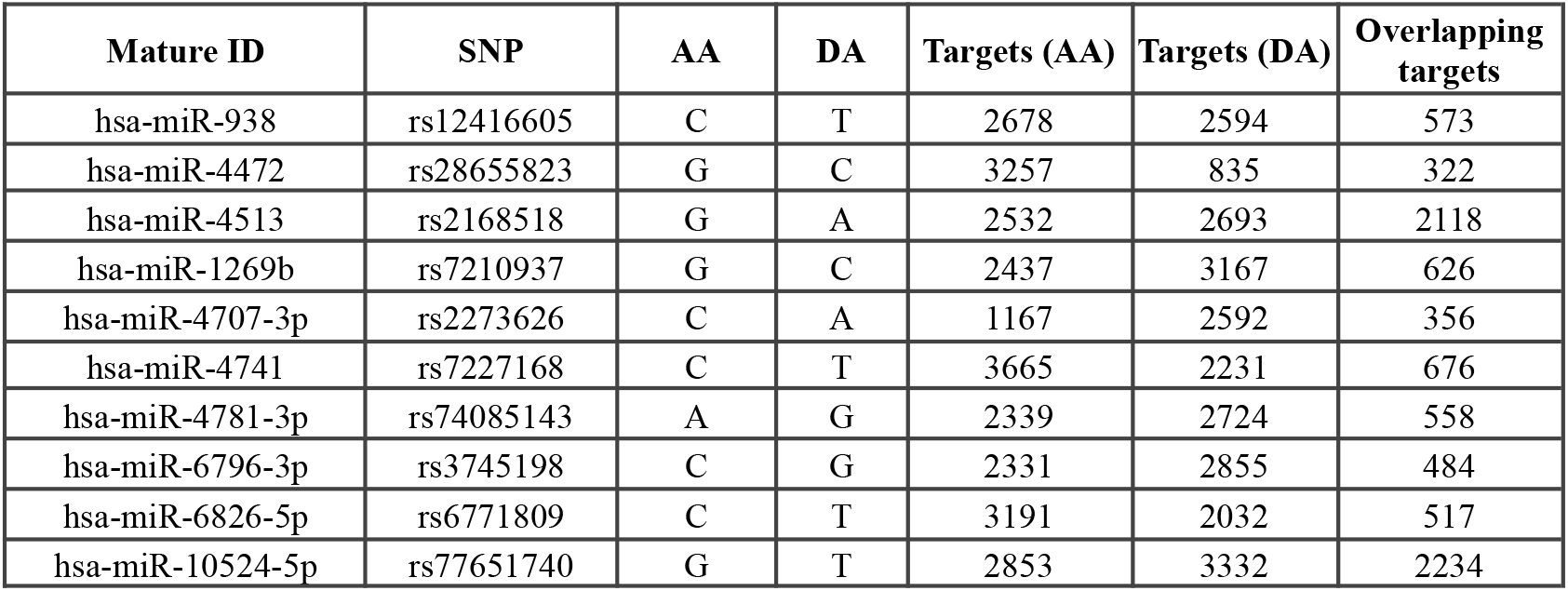
TargetScanHuman predicted target genes of the seed-variant miRNA candidates. Two sets of target genes were predicted for each candidate holding both ancestral (AA) and derived alleles (DA). The overlap between these two lists of target genes is provided and the similarity is estimated with the cosine similarity (see Methods)

| Mature ID | SNP | AA | DA | Targets (AA) | Targets (DA) | Overlapping targets |
| --- | --- | --- | --- | --- | --- | --- |
| hsa-miR-938 | rs12416605 | C | T | 2678 | 2594 | 573 |
| hsa-miR-4472 | rs28655823 | G | C | 3257 | 835 | 322 |
| hsa-miR-4513 | rs2168518 | G | A | 2532 | 2693 | 2118 |
| hsa-miR-1269b | rs7210937 | G | C | 2437 | 3167 | 626 |
| hsa-miR-4707-3p | rs2273626 | C | A | 1167 | 2592 | 356 |
| hsa-miR-4741 | rs7227168 | C | T | 3665 | 2231 | 676 |
| hsa-miR-4781-3p | rs74085143 | A | G | 2339 | 2724 | 558 |
| hsa-miR-6796-3p | rs3745198 | C | G | 2331 | 2855 | 484 |
| hsa-miR-6826-5p | rs6771809 | C | T | 3191 | 2032 | 517 |
| hsa-miR-10524-5p | rs77651740 | G | T | 2853 | 3332 | 2234 |

Next, we wanted to examine these candidate miRNAs with SNPs showing the highest population differentiation more in depth. We reviewed the literature looking for particular phenotypes in human populations and potential regulatory processes where these variants might be associated with. Among the ten miRNA candidates with SNPs located in the seed, all except one (hsa-miR-10524-5p) have been related to disease and, specially, with different types of cancers (Table 1), showing some of them differences among populations attributable to genetic risk factors, like in breast cancer (BC), colorectal cancer (CRC) and gastric cancer (GC) (Sung et al. 2021). Particularly, three of these miRNAs (hsa-miR-4472, hsa-miR-4513 and hsa-miR-6826-5p) were associated with BC, two (hsa-miR-4472 and hsa-miR-4741) with CRC and two (hsa-miR-938, hsa-miR-4513) with GC. In four out of the nine miRNAs related to disease the miRNA association was linked to the presence of the variant (rs12416605 in hsa-miR-938, rs7210937 in hsa-miR-1269b, rs2168518 in hsa-miR-4513 and rs2273626 in hsa-miR-4707-3p) (Table 1). When considering the 14 miRNAs candidates with SNPs located in the mature regions we observed that, all except one, for which no previous data have been reported (hsa-miR-6811), have been previously related to disease (Table 1). Among the associations with cancers showing differences on their risk among populations, five (hsa-miR-196a-3p, hsa-miR-646, hsa-miR-1269a, hsa-miR-6826-5p and hsa-miR-8084) have been associated with BC, five (hsa-miR-196a-3p, hsa-miR-646, hsa-miR-1269a, hsa-miR-6071 and hsa-miR-6826-5p) with CRC, and four (hsa-miR-196a-3p, hsa-miR-646, hsa-miR-1269a and hsa-miR-1304-3p) with GC. In four out of the 13 miRNAs related to disease the miRNAs association was linked to the presence of the variant (rs11614913 in hsa-miR-196a-3p, rs61992671 in hsa-miR-412-3p, rs6513497 in hsa-miR-646 and rs73239138 in hsa-miR-1269a) (Table 1).

In particular, for rs11614913 in hsa-miR-196a-3p (F_st_ = 0.24) the derived T allele has been associated with a decreased risk of different types of cancers, including breast and gastrointestinal cancers, principally in Asian populations. The frequency of the derived T allele is higher in East Asians (~ 54%) than in Europeans (CEU ~ 44%) and remarkably higher than in Africans (~13%) which may explain differences in the presentation of these types of cancer among populations and would agree with selective processes in this SNP. Similarly, for rs12416605 in hsa-miR-938 (F_st_ = 0.21), the derived T allele has been reported as a protective factor for the susceptibility to suffer a diffuse subtype of gastric cancer with the finding of a higher frequency of the T allele in Europeans compared with Asians (~29% vs. ~2%), which would agree with the reported higher predisposition to gastric cancer in Asian populations (Torruella-Loran et al. 2019). In this regard, also the T allele of rs73239138 in hsa-miR-1269a (F_st_ = 0.22) has been significantly associated with a decreased risk of gastric cancer in a chinese population (Table 1).

Although most of the literature is centered on cancer diseases, other pathologies showing population differences worldwide have been linked to some of these miRNA candidates and SNPs. The T allele of rs11614913 in hsa-miR-196a-3p (highest frequency in Asian populations: 54%) shows a pleiotropic effect being not only associated with cancer but also with the risk of developing coronary artery disease (CAD) (Fragoso et al. 2019), as well as the T allele of rs2168518 in hsa-miR-4513 (highest frequency in European populations: 61%), which has been strongly associated with increased susceptibility to CAD and other related pathologies and physiological states showing risk differences among populations such as glucose homeostasis, blood pressure, and age-related macular degeneration (Mir et al. 2019; Ghanbari et al. 2014 and 2017; Li et al. 2015).

Additionally, among the SNP candidates with the highest F_st_ scores in the top 1% is rs2273626 (F_st_ = 0.57), located in the seed region of hsa-miR-4707-3p. A neuroprotective role for the derived T allele in the progression of glaucoma has been reported (Ghanbari et al. 2017), which goes in line with the negative association of rs2273626 with the disease (Springelkamp et al. 2017). This SNP shows a derived allele frequency of ~3% in African populations and more than 50% in non-Africans (Fig. 4a), which would be in agreement with the higher incidence of glaucoma in Africans (Abu-Amero et al. 2015). Furthermore, the extended haplotype homozygosity (EHH) decay on this variant indicates the presence of longer haplotypes harbouring the derived allele in non-African populations (Fig. 4b), which is consistent with the occurrence of positive selection processes favouring the neuroprotective allele since the Out-of-Africa event.

**Fig. 4.**
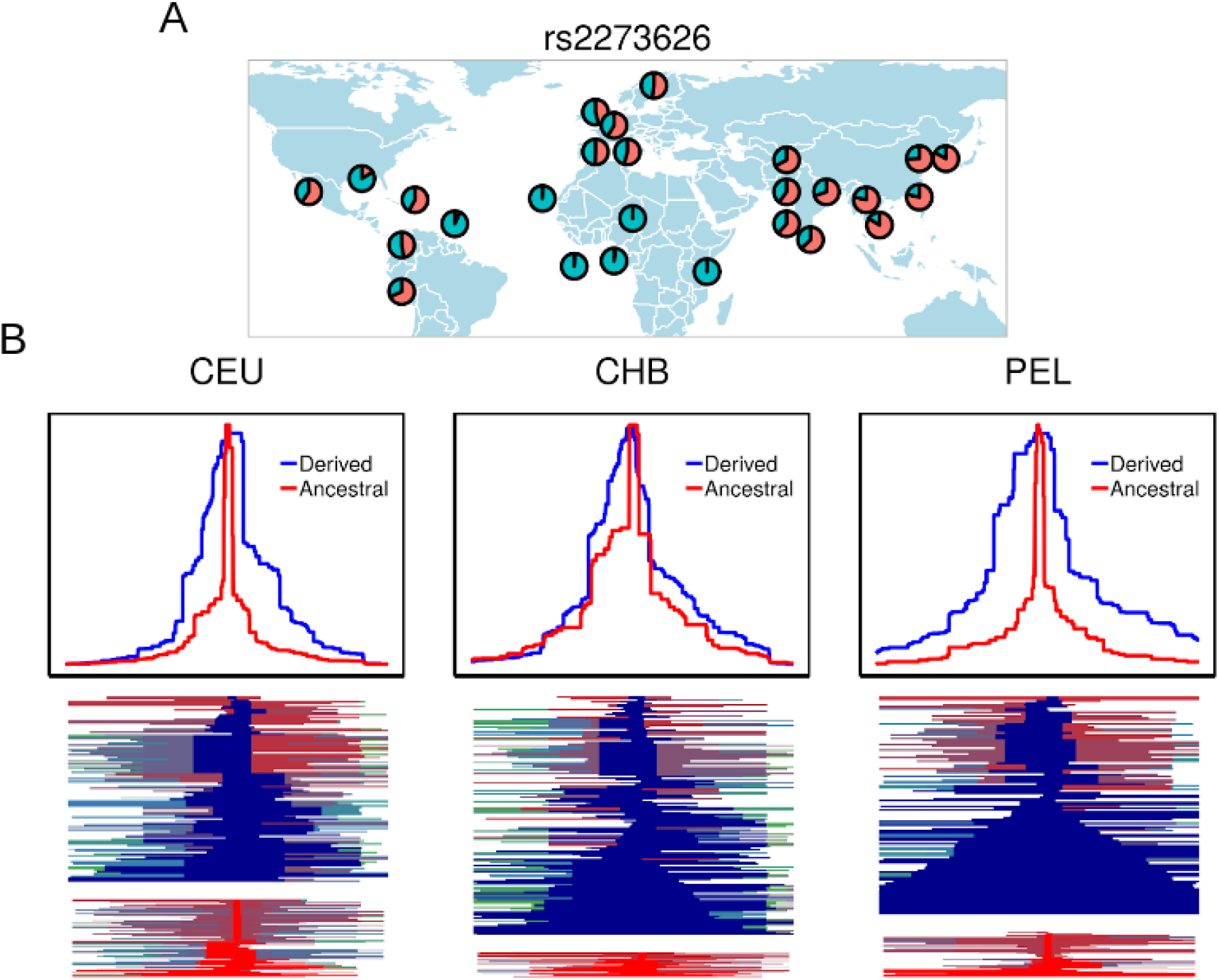
Analysis of signatures of positive selection in the candidate SNP rs2273626. **(a)** World wide MAF distribution of rs2273626. **(b)** Extended haplotype homozygosity (EHH) decay in both ancestral and derived alleles of rs2273626 (upper plot) and haplotype patterns around the ancestral and derived alleles (bottom plot) in Utah Europeans (CEU), Han Chinese (CHB) and Peruvian (PEL) populations

## Discussion

The increasing discovery of naturally occurring variation in the human genome, together with the improvement in annotation strategies of non-protein coding genes, has made it possible to study the potential consequences of mutations in the human miRNAs. As a dense layer of post-transcriptional regulation, miRNAs are expected to be highly susceptible to the occurrence of mutations in their sequences. However, in this analysis, along with previous studies (Carbonell et al. 2012), we discuss the unexpected level of variation in the critical regions of these regulatory molecules and its possible relationship with evolutionary processes associated with disease.

We implemented a computational pipeline to annotate and analyse the nucleotide diversity and selection signatures of the most updated catalog of genetic variation from 1000 Genomes Project (phase III), in the most complete collection of annotated human miRNAs to date (miRBase, v.22). We integrated the analysis of miRNA variation with the most sophisticated software for target prediction to date, *TargetScanHuman*, which was adapted to predict allele-specific target genes in seed-harbouring SNP miRNAs. This method, unlike others previously published (Riffo-Campos et al. 2016), incorporates multiple features from target conservation to sequence context to generate more accurate prediction scores. As a result, this provided a robust approach to compare the allele-driven targeting and estimate the extent of the shift generated in the gene target profiles of seed-harbouring SNP miRNAs. We also integrated novedous statistical methods sensitive to different modes of selective sweeps (hard and soft) to capture a wider range of selection signatures than previously reported for human miRNAs.

Until now very few studies have considered the integrated role of the different genomic factors that might have shaped the global diversity of the human microRNAome (Gallego et al. 2016). Here we show that the expansion of new miRNAs in the primate lineage, their location in the genome and the role of hosting transposable elements are significantly associated with the increase in miRNA diversity, something that might be related with the evolutionary boost of the miRNA system in the human genome. Furthermore, against the common belief, here we report a global excess of variation in the seed, which appears as the most diverse among the traditionally conserved functional regions of miRNAs. This is in contrast with the low diversity found in the loop, which evidences the evolutionary constraints due to its role in hairpin folding. This evidence stresses the importance of the secondary structure in maintaining the stability of the RNA molecule and determining the balance between miRNA biogenesis, particularly binding of the miRNA with the Drosha-DGCR8 complex, and miRNA turnover (Han et al. 2006; Guo et al. 2015). Moreover, the population differences found in these two regions are among the highest in the whole precursor sequence, something compatible with targeted evolutionary-driven processes that might be implicated in regulatory advantages. These processes are evaluated in the present study by identifying a global enrichment in positive selection signals (selective sweeps) among the highest differentiated SNPs across populations, showing the potential of these miRNAs and their regulatory networks to drive population-specific adaptations in agreement with some previously reported works (Quach et al., 2009; Li et al. 2012; Torruella-Loran et al. 2016).

Either by changing their targeting profiles or modifying their expression levels, it is clear that miRNA networks are more versatile to sequence changes than reported until now. We show that a significant fraction of human miRNAs participate in gene expression variation driven by the presence of eQTLs in their sequences. This goes in line with the regulatory plasticity that miRNAs have proven to hold and that might be determinant in adaptive changes at regulatory level. However, the phenotypic consequences of adaptive changes in these molecules are far to be properly understood. The great target breadth of miRNAs and the massive complexity of their regulatory networks make changes in their sequences affect multiple pathways simultaneously. Therefore, selective forces that rewire these networks might also be behind population-specific susceptibilities to different disorders. In this line, here we show that human miRNAs are also enriched in variants associated with specific human traits and diseases reported by GWAS studies. In this paper we provide a collection of miRNA alleles that were reported to affect individuals differently depending on their genetic ancestries.

In this regard, some of the miRNAs with SNPs showing the highest population differentiation have been found associated with diseases that show different population prevalence worldwide. One of the clearest examples is the case of rs12416605 in hsa-miR-938, whose derived T allele has been reported to confer protection against the diffuse subtype of gastric cancer (GC) through one of its targets, the chemokine *CXCL12* (Torruella-Loran et al. 2019), reported as playing a critical role in cell migration and invasion (Izumi et al. 2016). This cancer seems to be promoted by the amplified repression of *CXCL12,* mediated by the rs12416605 ancestral C allele (Torruella-Loran et al. 2019), which makes C-allele carriers more susceptible to develop GC metastasis. This would be in agreement with the finding of a higher frequency of the T allele in European compared with Asian populations, which is reflected by a high‐global fixation index (F_st_), and may influence the existing geographical clinical differences between Asian and non-Asian populations (Lin et al. 2015).

Among non-cancer diseases we found the T alleles of rs11614913 in hsa-miR-196a-3p and rs2168518 in hsa-miR-4513, associated with increased susceptibility to coronary artery disease (CAD). Although this disease seems to be highly dependent on environmental factors, with over 60% of current cases occurring in developing countries (Beltrame et al. 2012), population differences in CAD susceptibility are envisaged. In that context, the most striking finding is for primary open-angle glaucoma (POAG), a complex neurodegenerative disorder, dependent on environmental and genetic factors, that causes irreversible blindness and affects approximately 70 million people worldwide. Recent studies report a highly biased prevalence of the disease towards individuals with African ancestry, followed by Asians and Europeans (Abu-Amero et al. 2015). Several genes have been found associated with the progression of the disease by diverse GWAS studies. Among them, the caspase recruitment domain family member 10 (*CARD10*) seems to confer a neuroprotective role by increasing the survival and proliferation of retinal ganglion cells (Khor et al. 2011), whose apoptosis is enhanced in POAG. In Ghanbari et al. (2017), the authors demonstrated by allele-specific *in vitro* validation that the rs2273626 derived T-allele generates a lower repression of *CARD10*. A weaker binding to the target seems to be behind this expression change, which we further validated with *TargetScanHuman*, reporting a greater repression score by the ancestral allele (0.632) than the derived allele (0.124). The authors suggest that the neuroprotective role of *CARD10* in the progression of glaucoma is associated with this lower repression, supported by the negative association of rs2273626 with the disease (Springelkamp et al. 2017). Here we report that the allele-specific regulation of *CARD10* through hsa-miR-4707-3p might contribute to the ethnic disparities prevalence of POAG and that this differential regulation is driven by processes of positive selection that promote the neuroprotective role of rs2273626 derived T-allele in non-African populations.

Here we show that, despite the strong selective pressures that maintain miRNA conservation, several miRNA variants might have suffered the effect of positive selection and may account for phenotypic diversity among human populations being, in some cases, related to disease. Even though we identify some of these miRNA variants and, in certain cases, functional data shows allele-specific regulation of specific target genes, the extent to which most of these miRNA mutations contribute to differences in disease risk among populations remains to be investigated. One of the main limitations of the analysis of positive selection in miRNAs is their small size. Haplotype-based statistics like iHS and nSL rely on the detection of long unbroken haplotypes that might span thousands of base pairs on both sides of the selected locus, which hinder the identification of the true target of selection. The intronic origin of a substantial number of human miRNAs also makes difficult the identification of the causal genomic locus of the selection signature, potentially being originated either by the miRNA or the hosting gene. The conclusive evidence to understand the contribution of miRNAs to the recent evolutionary history of humans is the experimental validation of the genotype-phenotype association. However, the multiple potential targets of miRNAs and the side effects generated by sequence changes in the non-selected cellular processes makes this validation a difficult task. New methods and more data are needed to fill this gap between the genetic change and the phenotypic adaptation.

## Materials and Methods

### Human miRNA coordinates and functional region annotation

The human miRNA genomic coordinates were downloaded from the last release of the miRBase annotation database (v.22, March 2018) (Kozomara et al. 2019, http://www.mirbase.org/). This dataset contains the coordinates of 1918 human miRNA precursor transcripts and their mature sequences that were converted to hg19 genome assembly with liftOver (Hinrichs et al. 2006). From this conversion, four miRNA genes were dropped from the original dataset, and 10 were not able to be located in any chromosome, being also removed and leaving a total of 1904 precursor sequences. A custom script was designed to extract the individual functional regions of each miRNA. As shown in Fig. 1a we differentiated the “seed” region (positions 2-8), the mature (“mat”) region outside the seed, the “loop” (region between two mature sequences) and the precursor regions (5’ and 3’ sides) outside the mature and loop. We also considered precursor flanking regions on both sides (5’ and 3’) of each miRNA hairpin, having the same length as the whole precursor sequences. An additional category was created in order to accommodate the regions that overlap between different miRNAs (“ovlp”), these miRNAs are treated differently due to the difficulty of analysing the overlapping regions. In the analysis of region-specific diversity the miRNAs with “ovlp” regions (71) were discarded. A different degree of mature annotation is seen in the miRBase transcripts: 959 transcripts out of the 1904 (50.3%) present both mature sequences annotated (5p and 3p arms), allowing to completely describe the different regions of the precursor sequences. However, in 945 transcripts (49.7%) only one mature sequence is reported. In these cases, the description of the whole precursor sequence is limited to the boundaries of the single mature described (the specific boundaries of the loop region are not able to be defined). Therefore, when extracting the functional regions of the miRNA genes, the precursor region is considered as the whole portion that encompasses from the end of the given mature sequence to the start of the opposite flanking region (this would retain as “precursor” the “loop” region, the unannotated mature region and the actual premature region of that arm). The “loop” region is only extracted when the two mature sequence coordinates are given. These inconsistencies in the annotation of the miRNA transcripts are taken into account throughout the analysis (Fig. 1a).

### Computational analyses of genomic context, evolutionary age and clustering annotation

A computational pipeline was used to integrate the tools to annotate miRNAs, locate variants in the miRNA sequences and perform the statistical calculations for the analysis of diversity, positive selection and target prediction. This pipeline was adapted to work in a high performance computing (HPC) environment based on the cluster management and job scheduling system SLURM. In order to obtain the genomic context of miRNAs, we intersected the GENCODE 19 protein coding gene and lncRNA gene annotations (v.29) (Frankish et al. 2019) with the miRNA coordinates with the multipurpose software *Bedtools* (Quinlan et al. 2010), which allow us to find coordinate overlaps between two or more sets of genomic regions with a minimum overlap of 1bp (*Bedtools intersect* functionality). The RepeatMasker open-4.0.5 database (repeat library 20140131) (Smit et al. 2013-2015), which looks for interspersed repeats and low-complexity DNA (simple repeats, microsatelites), was also used in order to define the overlap of miRNAs with repetitive elements. miRNAs were classified based on their evolutionary age by merging the classifications obtained in Iwama et al. (2013) and Santpere et al. (2016). We grouped the miRNAs in the following categories: Primate-specific (group 1, previous 5 to 12 groups in Iwama et al. (2013)); Eutherians (group 2, previous 1 to 4); Metatheria and prototheria (group 3, previous −1 to 0) and Conserved beyond mammals (group 4, previous −2 to −3). The remaining 281 miRNAs were non-classified due to absence of data or discrepancies between the two studies in their evolutionary age. In order to obtain the miRNA clusters, a python-based custom script was designed to calculate the closest distance of each miRNA to any other in the same strand and chromosome. We defined miRNA clusters as groups of two or more miRNA genes separated by 10000 bp or less (Guo et al. 2014). The contributions of the genomic context, evolutionary age and clustering to the nucleotide diversity were obtained by applying a multiple linear regression model (*lm*), which is based on the programming language R (R Core Team 2020) and seeks to estimate the relationships between these factors (predictors) and the response variable (diversity).

### miRNA genetic variation and nucleotide diversity

Human variation data from The 1000 Genomes project (third phase) (Auton A et al. 2015) was used to annotate the human miRNA dataset. 26 different human populations accounting for a total of 2504 individuals were considered in the analysis, including the admixed populations from South Asia (SAS) and the Americas (AMR). We used the last version of the program *BCFtools* (v.1.11) (Danecek et al. 2021), for processing and analysing high-throughput sequencing data, to extract the variants located within the miRNA sequences. Only biallelic SNPs with a MAF greater or equal than 1% in individual populations and 0.5% in the global population were taken into account. In the case of unnamed variants, these were kept and corrected by using the physical position preceded by “rs_” as provisional SNP ID. When computing the derived allele frequency and haplotype-based statistics, the human ancestral alleles annotated in the original VCF files were used to format the REF and ALT fields and the corresponding genotypes of the individuals. Any SNP whose ancestral status was unknown or did not match with the reference or alternative alleles were removed from the dataset. The overall pairwise mismatches per SNP (pi) were calculated with *BCFtools* in the whole miRNA SNP dataset, after that the nucleotide diversity (Pi) per region was computed by obtaining the diversity per nucleotide in the whole length (L) of each functional region (Pi = pi/L). In this way we consider each category of region (flank, pre, mat, seed, loop) as a single sequence instead of calculating the nucleotide diversity in the regions of the individual miRNAs. The nucleotide diversity per position was calculated by aligning the precursor transcripts of the whole miRNA dataset and obtaining the mean pi value at each site. In this analysis, the “ovlp” regions were not taken into account due to the difficulty of interpreting the diversity properties of such overlaps.

### Pathogenicity and disease associations of miRNA variants

The catalog of Combined Annotation Dependent Depletion (CADD) scores (Rentzsch et al. 2019) provides a quantitative way to measure the deleteriousness of single nucleotide polymorphisms (SNPs) in the human genome by prioritizing the functionality and diseases causing variants. This catalog was used to assess the level of pathogenicity of miRNA-harbouring SNPs as aproxy of their functionality. According to Kircher et al. (2014) a threshold of PHRED-scaled CADD score ≥ 10 is normally used to discern the 1% most deleterious SNPs of the whole human genome. We also leveraged the GWAS (v1.0) catalog (Buniello et al. 2019) to evaluate the participation of miRNA-harbouring SNPs in human traits.

### Calculation of F_st_, iHS and nSL scores

Population fixation indexes (F_st_) were computed by using the Hudson estimator of the F_st_ statistic, which is not affected by the sample size and does not overestimate the F_st_ scores in comparison with others (Bhatia et al. 2013), in all the variant miRNAs. The calculations were performed by pairwise comparison between the 26 populations used from the 1000 Genome project dataset. These F_st_ scores were normalized by frequency by performing a linear regression of the estimator values and the global MAF, the residual values were used as the final F_st_ scores. We extended the analysis of selection with two haplotype-based statistics: iHS (Voight et al. 2006) and nSL (Ferrer-Admetlla et al. 2014). These tests rely on the detection of blocks of homozygosity by the EHH statistic (Extended Haplotype Homozygosity) introduced by (Sabeti et al. 2002). A recent positive selection signal is found when these blocks present moderately high or intermediate frequency of derived alleles. The iHS test is designed to detect ongoing hard sweep signals, signatures characterized by the presence of a single sweeping haplotype at high frequency in their way to fixation. On the other hand, nSL was designed to detect either ongoing hard and soft sweep signatures with a greater power than iHS. In the case of soft sweeps, these are signatures of selection on standing variation, where more than one haplotype is sweeping at intermediate frequencies. The calculations of iHS and nSL were computed with the software *selscan* (Szpiech et al. 2014), an application that implements different haplotype-based statistics in a multithreaded framework. We allowed for a maximum gap of 20kb and kept only SNPs with a minor allele frequency (MAF) higher than 5%. This statistic is standardized (mean 0, variance 1) by the distribution of observed scores over a range of SNPs with similar derived allele frequencies. The standardization was performed in each population separately by using the *norm* function, also contained in the *selscan* package (Voight et al. 2006).

### Target predictions

The program TargetScanHuman (TSH, release 7.2) (Agarwal et al. 2015) was used to perform the miRNA target predictions. The perl-based pipeline used by the authors (http://www.targetscan.org/cgi-bin/targetscan/data_download.vert72.cgi), together with the ViennaRNA package (Lorenz et al. 2011), were implemented locally and adapted to our needs of performing predictions from a custom miRNA dataset. This pipeline is composed by three different steps: (i) target site identification across the set of 3’UTR regions of the human genome, (ii) the probability of conserved targeting (P_ct_) calculations and (iii) the calculations of the context++ scores, which integrates different genomic features implicated in targeting efficiency. miRNA families and species information were downloaded from the *targetscan.org* Data Download page. In order to calculate the P_ct_ parameters, the 3’UTR dataset from the GENCODE version 19 (Ensembl 75) was obtained as a 84-way alignment from the same download page. As described in Agarwal et al. (2015), only the longest 3’UTR isoform of each gene was used as representative transcripts. In order to account for the miRNA variation in the target predictions, the variable positions in the miRNA seed regions (ancestral and derived states) were considered and incorporated into the TSH pipeline. Two different miRNA datasets were obtained when accounting for the ancestral and derived alleles of the SNPs found in the seed regions. As described in Agarwal et al. (2015), the accumulated weighted-scores per target gene were calculated as the sum of the individual target site weighted-scores, which is the final score associated with each target gene. As suggested by the authors, in order to remove the potential false positives we applied a custom per-site-based filtering strategy. Since negative weighted scores are associated with mRNA repression, only the per-site weighted scores below zero are considered and, from these, the per-miRNA 50th percentile was used as threshold to obtain the putative true target sites in each miRNA. In order to analyse the overlap between the predicted targets of the derived and ancestral miRNA alleles we used the cosine similarity (Hill et al. 2014), which is calculated by the total number of overlapping genes divided by the square root of the product of the number of targets of both alleles.

### Analysis of expression levels and expression variation

The catalogue of expression Quantitative Trait Loci (eQTLs) provided by the Genotype-Tissue Expression (GTEx) Project (Aguet F et al. 2017) was used to assess the implication of miRNA-harbouring variants in expression variation. Expression data from 16 different human tissues (bladder, blood, brain, breast, hair follicle, liver, lung, nasopharynx, pancreas, placenta, plasma, saliva, semen, serum, sperm and testis) was taken from Panwar et al. (2017). We used 2085 mature miRNAs from this dataset for which evolutionary age was available. Reads per million (RPM) values were analyzed for each mature miRNA separately, whose conservation status were determined by the precursor molecule following the classification criteria described before. A miRNA was considered to be expressed in a specific tissue when its reads were unequal to zero in at least one sample from that tissue. For the comparative analyses of the expression levels among conservation groups we took the total number of reads in the 16 tissues for all the miRNAs within each group.

## Acknowledgments

This study has been possible thanks to grant PID2019-110933GB-I00/AEI/10.13039/501100011033 awarded by the Spanish Agencia Estatal de Investigación (AEI), Ministerio de Ciencia, Innovación y Universidades (MCIU, Spain), the support of Secretaria d’Universitats i Recerca del Departament d’Economia i Coneixement de la Generalitat de Catalunya (GRC 2017 SGR 702), part of the “Unidad de Excelencia María de Maeztu”, funded by the AEI (CEX2018-000792-M), the Chilean Agencia Nacional de Investigación y Desarrollo - ANID (FONDECYT regular nº 1170446) and the Chilean Ministry of Education (MAG-1995). P.V-M is supported by an FPI PhD fellowship (FPI-BES-2016-077706) part of the “Unidad de Excelencia María de Maeztu” funded by MINECO (ref: MDM-2014-0370).

## Authors’ contributions

YE-P, HL conceived the study. PV-M, AG, HL and YE-P analysed and interpreted the data. PV-M, AG and YE-P wrote the manuscript. PV-M, AG, HL, YE-P and JB revised the manuscript. All authors approved the final manuscript.

## SUPPLEMENTARY MATERIAL

*See supplementary figures and tables in separated files. Titles and captions are provided below.*

**Supplementary File 1**: File including supplementary figures. **Fig. S1**: Number of annotated miRNAs in miRBase and their completeness. **Fig. S2**: (a) Frequencies of transposable elements whole genome and hosting miRNAs. (b) Chi square residuals of the association between genomic context categories and evolutionary ages of miRNAs. (c) Cumulative frequencies of distances between closely located miRNAs in the human genome. **Fig. S3**: Genomic location of human miRNA clusters. **Fig. S4**: (a) Nucleotide diversity (Pi) distributions of transposable elements hosting miRNAs. (b) SNP density in human miRNA regions. (c) Mean nucleotide diversity (Pi) of miRNA regions across the whole range of SNP MAF. **Fig. S5**: Mean nucleotide diversity (Pi) of miRNA seed regions in the 26 analysed populations from 1000 Genomes Project.

**Supplementary File 2**: File including supplementary tables. **Table S1**: Annotation of the human miRNA dataset (miRBase, v.22). **Table S2**: Number of miRNAs hosted by the different transposable element families. **Table S3**: Number of miRNAs found in the different genomic context categories.

