## Supplementary File 1 for "Signatures of genetic variation in human microRNAs point to processes of positive selection related to population-specific disease risks"

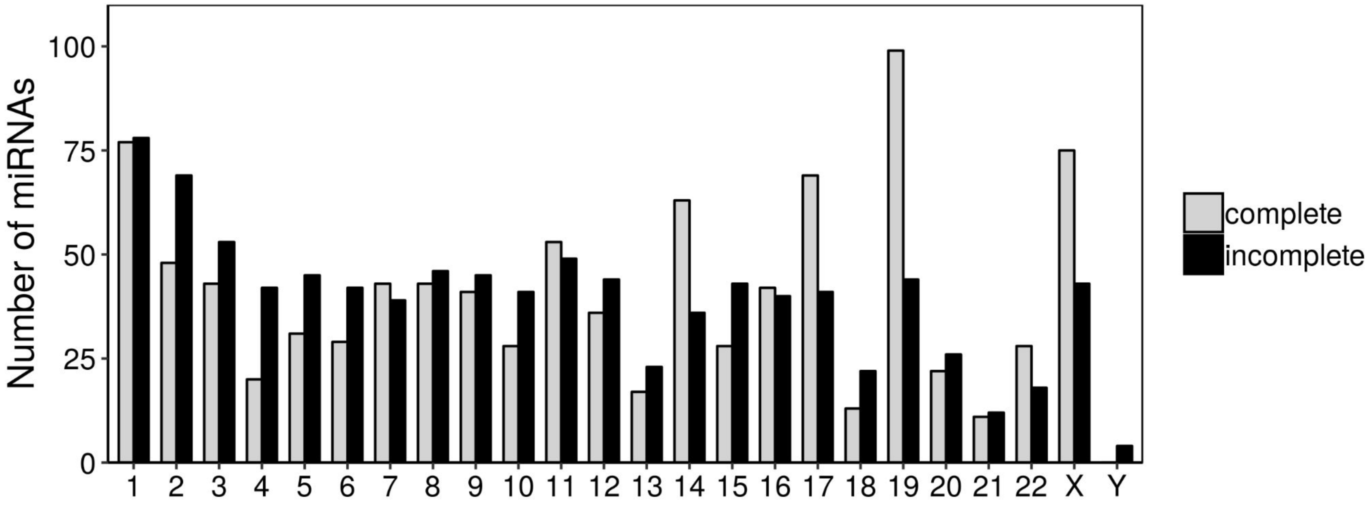


**Supplementary Fig. S1** Number of miRNAs per chromosome that present both mature sequences in their hairpin (complete annotation) and only one mature sequence in one of their arms (incomplete annotation)

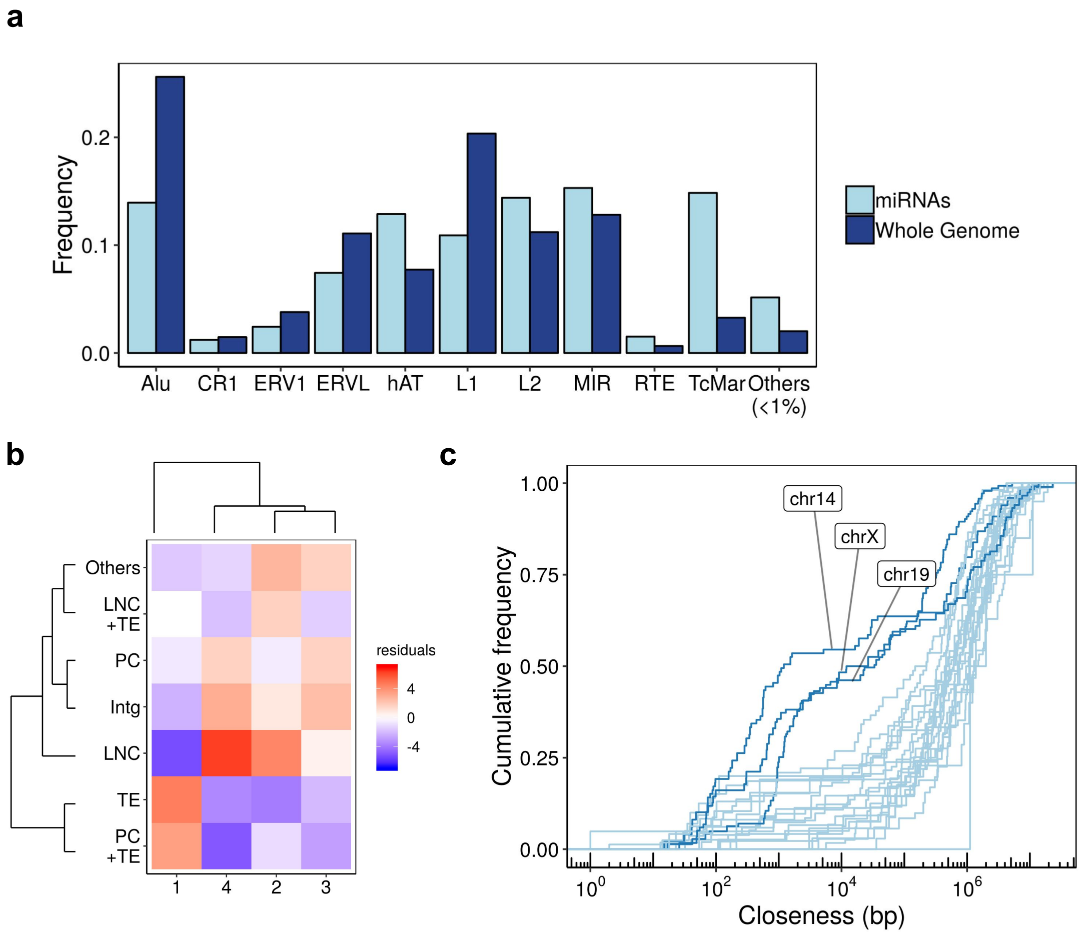


**Supplementary Fig. S2 (a)** Frequencies of transposable elements described by the RepeatMasker database (v4.0.5) in the whole genome and found overlapping miRNA sequences. **(b)** Chi square residuals associated with each of the genomic context categories across conservation groups. Dendrograms show the hierarchical clustering performed across rows (genomic context) and columns (conservation). **(c)** Cumulative frequency of the closeness found between miRNAs (distance to the closest miRNA) in each chromosome. The increase of frequency in chromosomes 14, 19 and X show groups of highly close miRNAs that correspond to the main clustering hotspots in the human genome

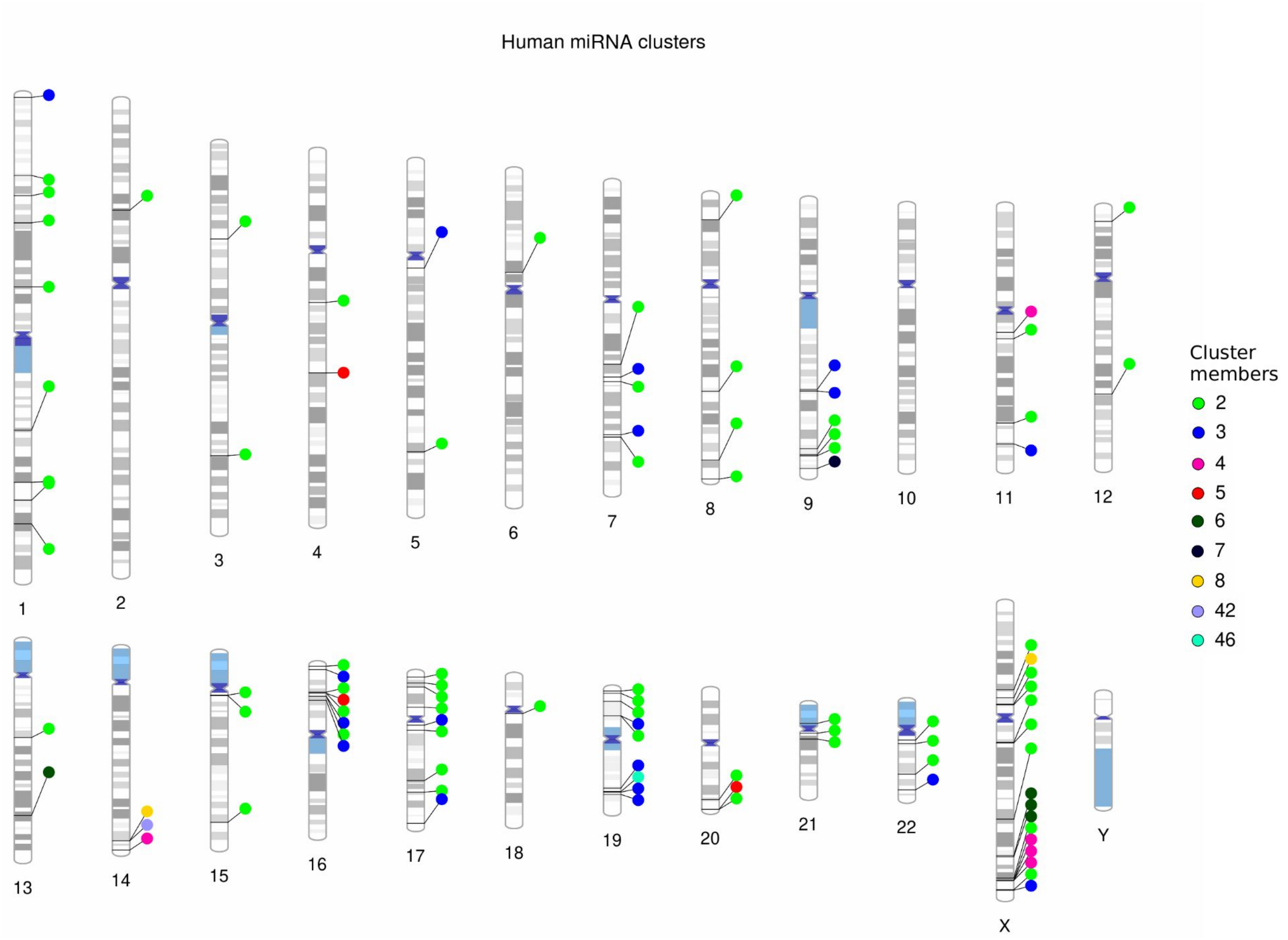


**Supplementary Fig. S3** Genomic location of human miRNA clusters

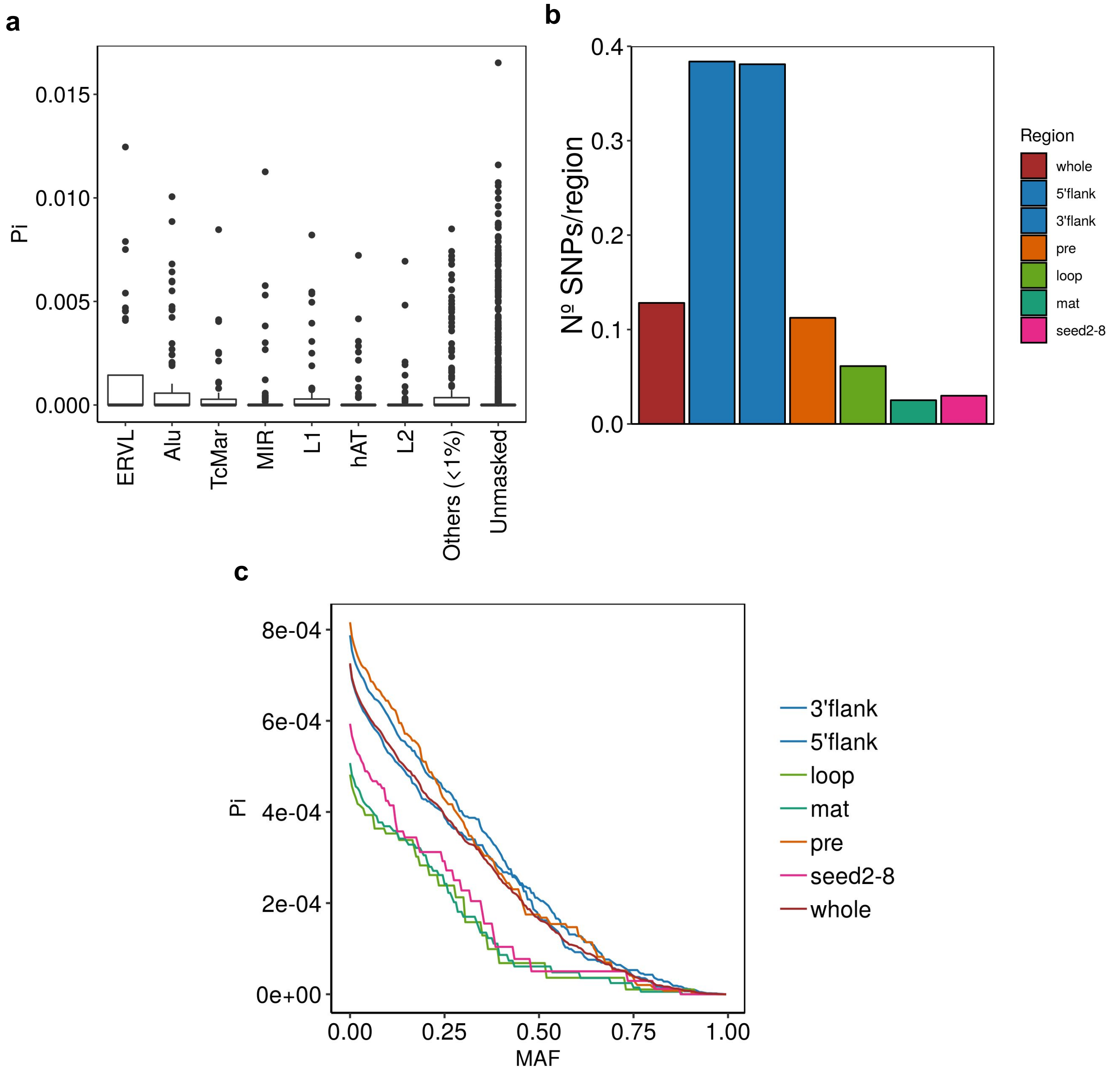


**Supplementary Fig. S4 (a)** Nucleotide diversity of miRNAs hosted by the different families of transposable elements. The “Others” category is made by minor categories represented by less than 1% of the total miRNAs. **(b)** SNP density per functional region calculated in the whole miRNA dataset. **(c)** Mean nucleotide diversity of the miRNA functional regions across the SNP MAF range

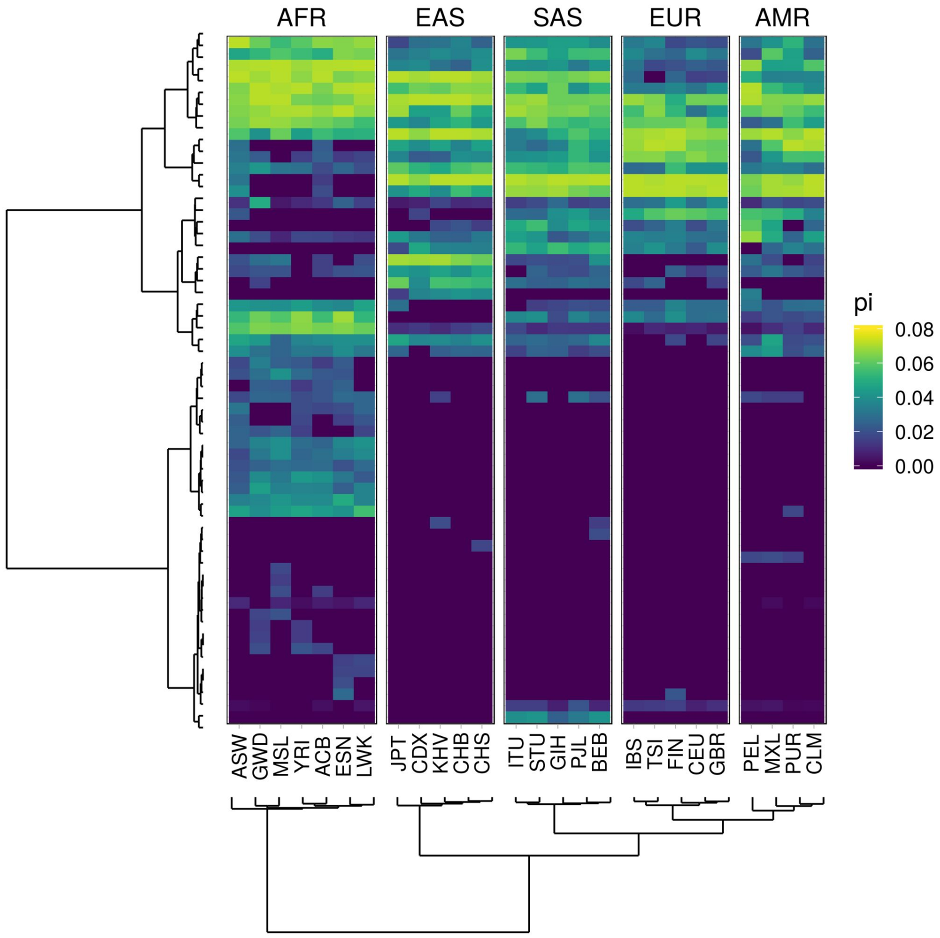


**Supplementary Fig. S5** Heatmap showing the mean nucleotide diversity values per population of the seed regions harbouring one or more SNPs of the whole dataset. The dendrograms represent the hierarchical clustering performed on the miRNAs (rows) and populations (columns)
